# Seipin transmembrane segments critically function in triglyceride nucleation and lipid droplet budding from the membrane

**DOI:** 10.1101/2021.12.05.471300

**Authors:** Siyoung Kim, Jeeyun Chung, Henning Arlt, Alexander J. Pak, Robert V. Farese, Tobias C. Walther, Gregory A. Voth

**Author notes:** Corresponding Author: Gregory A. Voth.

## Abstract

Lipid droplets (LDs) are organelles formed in the endoplasmic reticulum (ER) to store triacylglycerol (TG) and sterol esters. The ER protein seipin is key for LD biogenesis. Seipin forms a cage-like structure, with each seipin monomer containing a conserved hydrophobic helix (HH) and two transmembrane (TM) segments. How the different parts of seipin function in TG nucleation and LD budding is poorly understood. Here, we utilized molecular dynamics simulations of human seipin, along with cell-based experiments, to study seipin’s functions in protein-lipid interactions, lipid diffusion, and LD maturation. All-atom (AA) simulations indicate that most seipin TM segment residues located in the phospholipid (PL) tail region of the bilayer attract TG. We also find seipin TM segments control lipid diffusion and permeation into the protein complex. Simulating larger, growing LDs with coarse-grained (CG) models, we find that the seipin TM segments form a constricted neck structure to facilitate conversion of a flat oil lens into a budding LD. Using cell experiments and simulations, we also show that conserved, positively charged residues at the end of seipin’s TM segments affect LD maturation. We propose a model in which seipin TM segments critically function in TG nucleation and LD growth.

## INTRODUCTION

The lipid droplet (LD) is a fat-storing organelle, surrounded by numerous coat proteins and a phospholipid (PL) monolayer (Thiam et al., 2013; Walther et al., 2017). LDs store excess metabolic energy as highly reduced carbon triacylglycerol (TG) and can mobilize fatty acids for energy generation or membrane biosynthesis (Ducharme and Bickel, 2008; Walther and Farese, 2012). Due to their key role in metabolism, failure of LD biogenesis leads to metabolic diseases, such as lipodystrophy. Additionally, overwhelming the capacity of cells to form LDs is thought to be crucial for the development of diseases linked to obesity, such as fatty liver disease (Greenberg et al., 2011).

Current models of LD biogenesis posit that lipid droplet assembly complexes (LDACs) in the endoplasmic reticulum (ER) bilayer determine LD formation sites and facilitate LD growth (Arlt et al., 2021; Chung et al., 2019; Prasanna et al., 2021). LDACs, consisting of seipin and lipid droplet assembly factor 1 (LDAF1) in humans, or seipin/Fld1, Ldb16, and Ldo in yeast, efficiently catalyze the initial stages of LD formation (Chung et al., 2019; Klug et al., 2021; Teixeira et al., 2018; Wang et al., 2014). Absence of seipin, effectively removing LDAF1 as well (Chung et al., 2019), changes LD number and morphology dramatically, resulting in aggregated, small LDs and/or few supersized LDs (Fei et al., 2008; Salo et al., 2016; Szymanski et al., 2007; Wang et al., 2016). Therefore, investigating how seipin works is key to understanding LD biogenesis.

Human seipin is an undecamer, forming a ring-structure in the ER. Each subunit contains a lumenal domain, flanked by two transmembrane (TM) segments and short cytoplasmic tails (Arlt et al., 2021; Chung et al., 2019; Lundin et al., 2006; Sui et al., 2018; Yan et al., 2018). The lumenal domain has a conserved hydrophobic helix (HH) thought to insert into the lumenal leaflet of the ER membrane. It was suggested that the HH of human seipin, and in particular S165 and S166, is a key tethering site for TG (Prasanna et al., 2021; Zoni et al., 2021b) and might provide a binding site of LDAF1 (Chung et al., 2019). Yeast seipin lacks the HH, which may explain why it is not sufficient for function in LD formation (Arlt et al., 2021; Klug et al., 2021; Wang et al., 2014). A recent study on yeast seipin suggests its binding partner, Ldb16, provides several serine and threonine residues in the PL tail region, thereby working as a replacement of the conserved HH of seipin (Klug et al., 2021). It was also reported that seipin TM segments are required for seipin function (Arlt et al., 2021; Chung et al., 2019). Chimeric seipin proteins in which two TM segments were replaced with that of another ER protein (FIT2), were not functional and unable to form LDs while they can form a seipin oligomer (Arlt et al., 2021; Becuwe et al., 2020; Chung et al., 2019). This implies a crucial function for the evolutionarily conserved TM segments in LD biogenesis.

In this study, we capitalized on new information on seipin TM segments to investigate their roles in TG nucleation and ER-LD bridge formation using all-atom (AA) and coarse-grained (CG) molecular dynamics (MD) simulations. We discover a cage-like geometry of seipin TM segments facilitates a conversion of a planar oil lens into a unique ER-LD neck structure. Using cell experiments and CG simulations, we provide evidence that conserved, positively charged residues at the end of seipin’s TM segments are critical for LD maturation.

## RESULTS

Seipin TM segments are thought to be critical for seipin functions (Chung et al., 2019). The resolved structures, however, do not include TM segments likely because of their flexibility (Chung et al., 2019; Yan et al., 2018). Therefore, we modeled seipin structure including the TM segments with the residues ranging from Arg23 to Arg265 (Fig. 1) based on a yeast seipin structural model (Arlt et al., 2021) and our cryoelectron microscopy data of purified human seipin that partially resolved the TM segments (Fig. S1 of Supporting Information; see also Methods).

**Figure 1.**
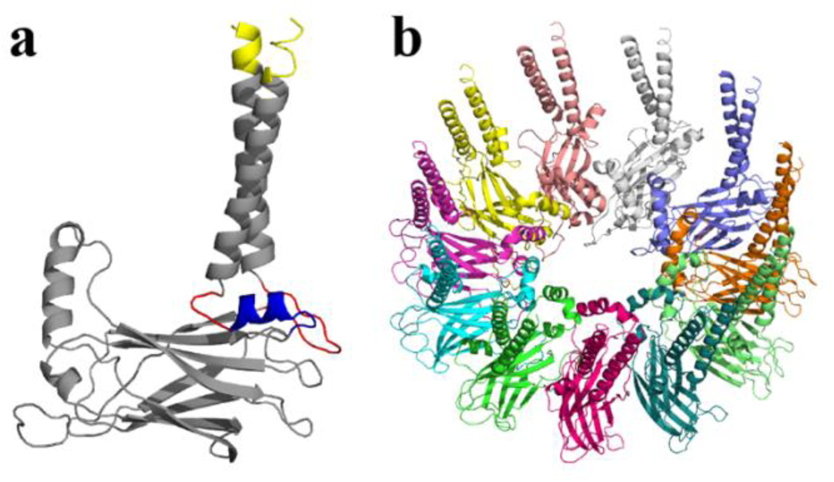
Structural model of human seipin. (a) Structure of a seipin subunit. The structure included in the cryoelectron microscopy data is shown in gray. Red loops were modelled with Modeller (Fiser et al., 2000). The blue region was predicted using the yeast structure. The helical structures were extended (yellow). (b) Structure of the human seipin oligomer used in the simulations. Each chain is shown with different colors.

How each part of seipin functions in LD biogenesis is not clearly known. To analyze the interactions between protein residues and lipids, we performed the AA simulation of human seipin in a 3-palmitoyl-2-oleoyl-D-glycero-1-phosphatidylcholine (POPC) bilayer containing 6% TG for 3 µs. To study protein-lipid interactions, we reduced the resolution of the AA simulation by molecularly grouping each lipid or protein residue as illustrated in Figs. 2a and 2b. We then calculated the normalized coordination number by molecule or the coordination number per molecule (||*S*||). This quantity, thought of as the concentration-independent coordination number, indicates how much each protein residue prefers interaction with PL or TG (see Methods and Fig. 2c). The HH exhibited a narrow spike, indicating preferential aggregation with TG (Fig. 2d). S166 had the largest value with TG in the analysis, consistent with other computational studies using CG simulations with the Shinoda-DeVane-Klein (SDK) or MARTINI force fields (Prasanna et al., 2021; Zoni et al., 2021b). Although the modified parameters of TG have a reduced charge distribution to reproduce the interfacial tension against water (Kim and Voth, 2021), the TG glycerol moiety can form hydrophilic interactions with protein residues in our AA simulation (e.g. S166). In contrast, the N- and C-terminal TM segments weaker but broader interactions with TG. Because N- and C-terminal TM segments, and HH, have helical structures, the attraction map had a weakly-defined periodicity (Fig. 2d). For instance, V163, S166, and F170 faced the membrane center, increasing the accessibility of TG. In contrast, F164 and L168 faced the lumenal interface, which prevented interactions with TG.

**Figure 2.**
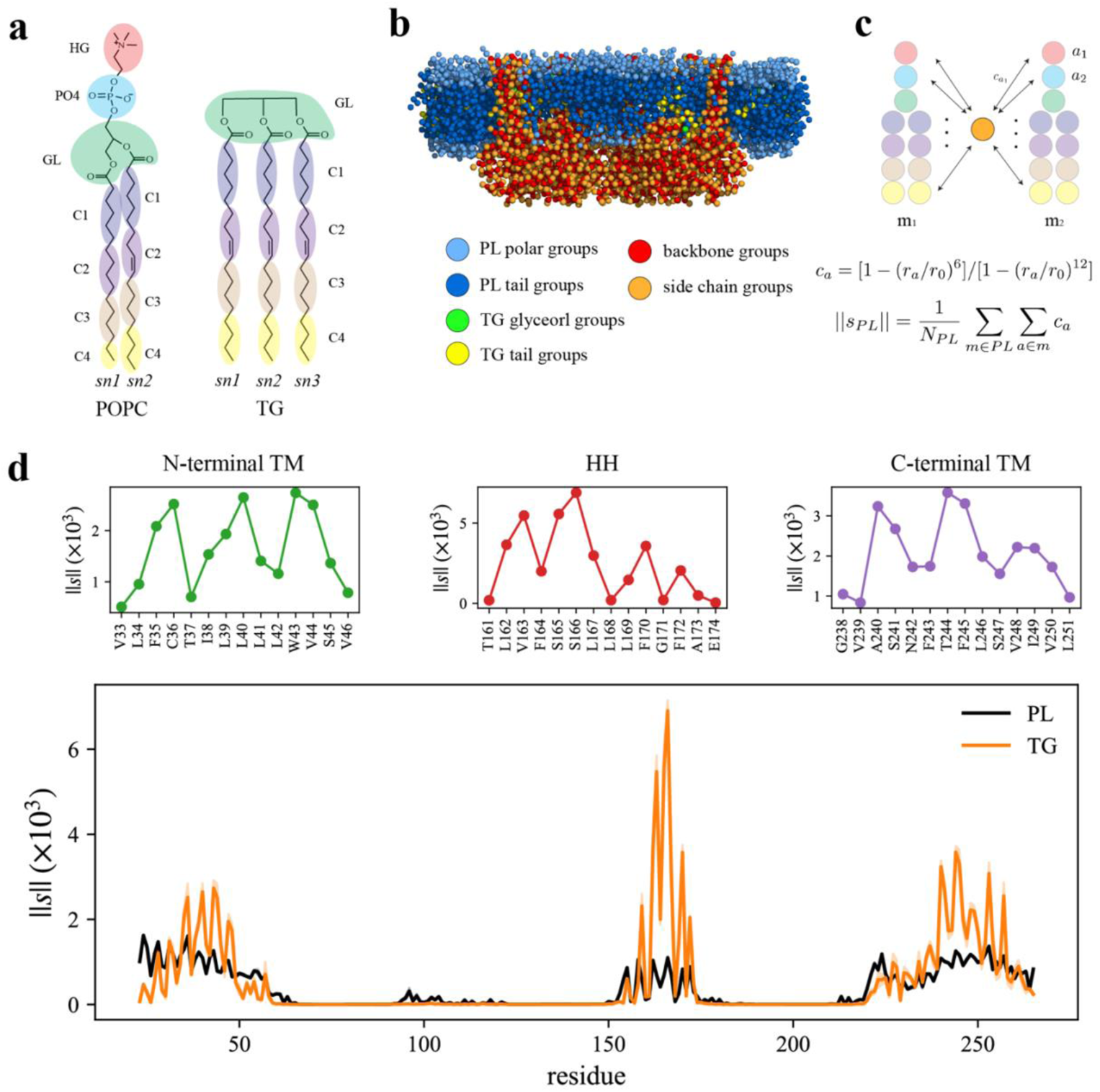
Seipin HH and TM segments attract TG. (a) Molecular groupings of lipids. Each protein residue was mapped onto one side chain and one backbone molecular group. (b) Initial structure of the system at the reduced resolution. The snapshot was clipped in the XZ plane. (c) Illustration of the calculation of the coordination number per molecule. A side chain group was depicted with orange circle and PL groups with other colors. (d) Interaction plot of protein residues with PL (black) and TG (orange). The shaded area represents the standard error of the results of three equal-length blocks, each containing 1-µs AA MD trajectory. The residues that had high interactions with TG in the N-terminal segment, HH, and C-terminal segment were shown in separate plots in the upper panel, colored with green, red, and purple lines, respectively.

We further compared protein residues’ attractions to TG glycerol groups or TG tail groups by normalizing the coordination number by the number of CG atoms. While the coordination number per molecule ((||*S*||)) indicates a propensity of each protein residue for each lipid type, PL or TG (Figs. 2c and 2d), the coordination number per CG atom (||*S*_*A*_||) provides a propensity for each CG atom type, in this case, a TG glycerol group or TG tail group (Fig. S2). As expected, S166 had a strong interaction with TG glycerol groups as they form a hydrophilic interaction (Fig. S2). The alignment of the insertion depths of S166 and TG glycerol moiety likely amplified the interaction. W257, which can form a hydrophilic interaction with TG glycerol groups, had a high value as well (Fig. S2). However, we note that those results were normalized by the number of CG atoms. If we compare the coordination number of TG glycerol groups and that of TG tail groups, TG tail groups will mostly have a higher value because there are 12 hydrophobic tail groups and 1 glycerol group for each TG molecule at the reduced resolution (Fig. 2a). Therefore, while hydrophilic interactions at the hydrophobic phase are significant, the largest driving force of TG nucleation inside the seipin ring is provided by hydrophobic interactions of TG with seipin HH and N- and C-terminal TM segments.

The architecture of the seipin complex results in an unusual ring of TM helices, which may prevent the exchange of molecules between its interior and exterior. To study how molecules permeate through the dense TM region of the seipin ring, we analyzed the orientation of TM segments. The order parameter, *S* (see Fig. 3a), was computed from the angle between each TM segment and the lower plane of the beta-sandwich region. If *S* is equal to 1, it represents the orientation of the TM segment vertical to the plane, and if *S* is −0.5, it is parallel to the plane. We also measured the distance (*d*) between the center of the masses of the N- and C-terminal TM segments. Both N- and C-terminal TM segments showed a broad range of orientations (Fig. 3a). Especially, the N-terminal TM segment sampled a broader range of angles than the C-terminal TM segment, likely because the switch region (Phe220-Phe230) which connects the lumenal domain with the C-terminal TM (see Figure 1a, blue) added some restraints on the movement. High flexibility of the TM segments was also supported by the calculation of the root-mean-square distance (RMSD) and root-mean-square fluctuation (RMSF) of α-carbon atoms, shown in Figs. S3 and S4, respectively. When computing the RMSD of the α-carbon atoms of the lumenal domain of a single subunit, it leveled off to 0.2 nm. However, when computing the RMSD of the whole subunit, it had higher values with larger error bars. Finally, the calculated RMSF showed high flexibility of both N- and C-terminal TM segments. The switch region itself was flexible but restrained the movement of the C-terminal TM close to it.

**Figure 3.**
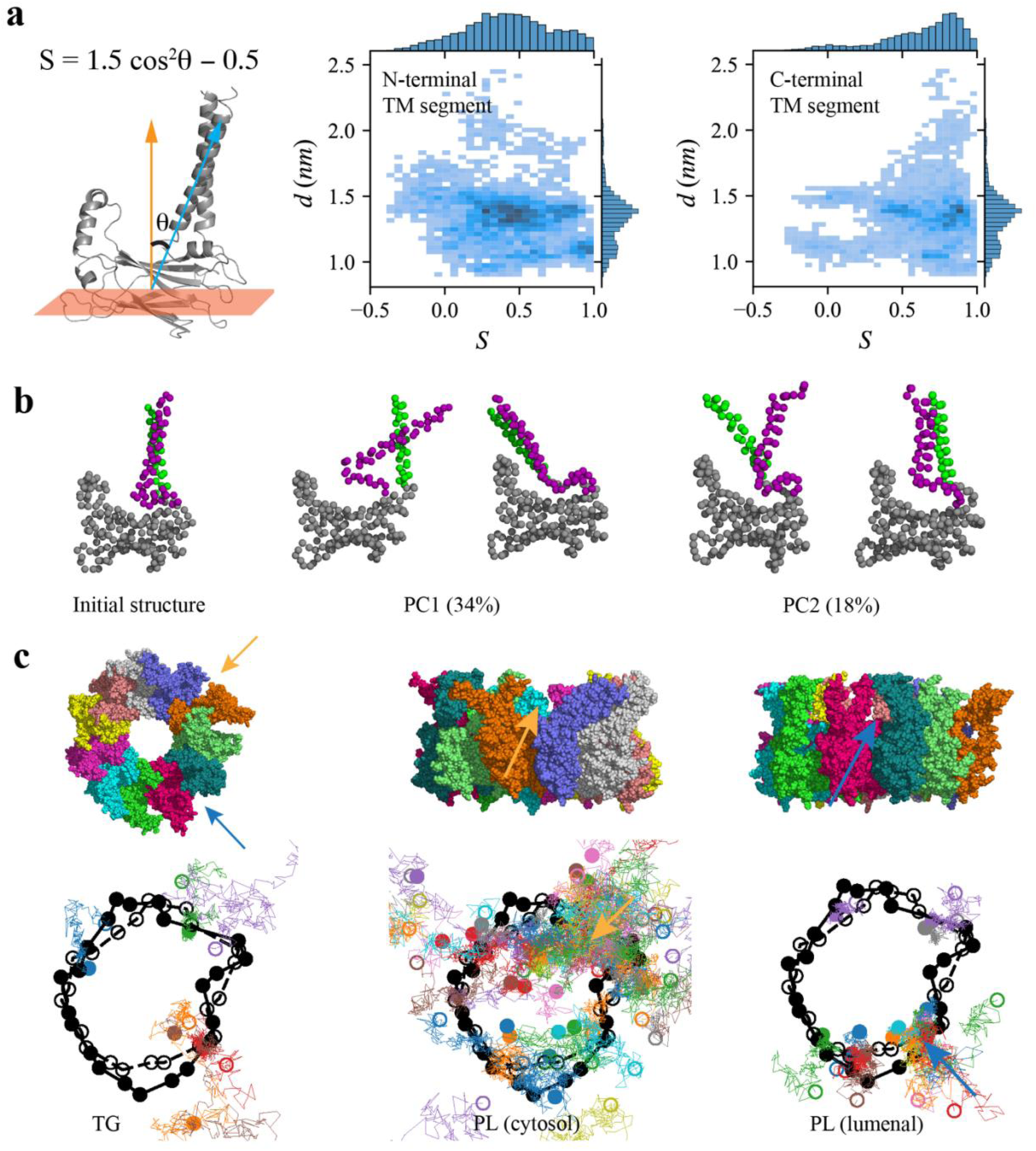
Flexibility of seipin TM segments increases lipid permeability. (a) Illustration of the angle (θ) of N- or C-terminal TM segments with the lower beta-sandwich plane and distance (*d*) between the center of masses of N- and C-terminal segments. (b) Two principal collective motions with explained variances of 34% and 18%, respectively. (c) Snapshots of the last frame in top and side view (top). The XY positions of TG, cytosolic PL, and lumenal PL that permeate through the TM region are shown every 5 ns with colored lines (bottom). Black markers indicate the XY positions of N- and C-terminal TM segments. Filled circles indicate the positions in the last frame and unfilled circles in the first frame.

To understand the structural changes of seipin, we aligned each subunit trajectory against its average structure and performed the principal component analysis (PCA) (Pearson, 1901) using the coordinates of α-carbon atoms. Because the TM segments were most flexible in seipin, the dominant collective motions were related to the movements of seipin TM segments (Fig. 3b). For instance, the first principal motion with an explained variance of 34% was a swing-back-and-forth motion of N- and C-terminal TM segments together. The second principal motion with an explained variance of 18% was sliding of N- and C-terminal TM segments into the opposite directions. Interestingly, we observed that fluctuations and flexibility can open the space between monomers, significantly increasing lipid permeation through the protein-dense TM region (Fig. 3c). For instance, the orange monomer and purple monomer in the snapshots of Fig. 3c open to generate space in the cytosolic leaflet between them, which caused increased lipid influx (orange arrow). Similarly, the pink and dark green monomers had dispersed TM segments in the lumenal leaflet, promoting lipid permeation (blue arrow). As TG is located closer to the membrane center than PL, TG permeation benefits from opening of TM segments in either the cytosolic leaflet or lumenal leaflet of the membrane. We also found the lumenal leaflet had more limited PL permeability than the cytosolic leaflet because of the switch region. Thus, while seipin has a dense array of TM segments, their flexibility increases permeability of lipids in and out of the complex. To understand how seipin influences the dynamics of lipids, we computed the position-dependent diffusion coefficient relative to the center of the mass of the lumenal domain (Fig. 4). While all lipids had comparable diffusion coefficients in the protein-free region (7.5 nm - 10.0 nm away from the seipin center), diffusion became slower near the TM segments and HH due to interactions with the protein. The slower diffusion near the TM region can also be seen in the lipid trajectories where the displacement became smaller near the TM segments (Fig. 3c). The decreased rate of diffusion coefficient is correlated with the contact area of protein. For instance, lumenal PLs up to 7 nm from the seipin center had the lowest diffusion coefficients because of the HH and switch region in the lumenal leaflet. In contrast, the cytosolic leaflet only contained the N- and C-terminal TM segments at the seipin boundary, leading to higher diffusion coefficients. The diffusion coefficients for TG were between those of cytosolic and lumenal PLs because TG molecules close to the lumenal leaflet can interact with the HH and switch region. In addition, strong attractions of TG with protein residues can further reduce the rate of diffusion (Fig. 2d). Due to confinement, the mean squared displacement (MSD) of the lumenal PLs trapped inside the seipin ring, referred to as proteinized PLs, leveled off at later simulation times (Fig. S5). Such confinement can increase the bending modulus of this area (Schachter et al., 2020), thereby working as a rigid base to ensure the direction of LD budding to the cytosolic side along with the lumenal domain of seipin.

**Figure 4.**
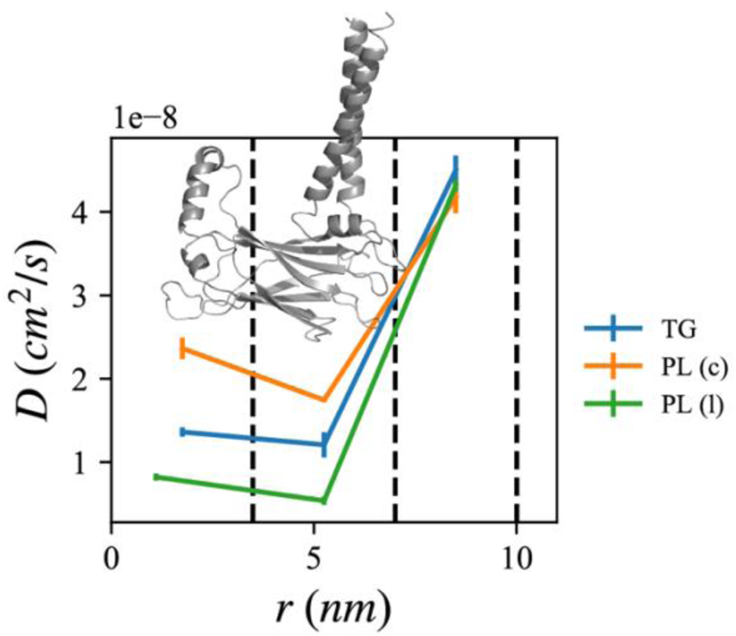
Lipid diffusion rate depends on location relative to the seipin oligomer. The center of the mass of the lumenal domain of seipin is at origin. The first region (0-3.5 nm) contains the seipin hydrophobic helices and the second region (3.5 nm - 7.0 nm) the TM segments. The third region (7.0-10.0 nm) is protein-free. Diffusion coefficients of TG, cytosolic PL, and lumenal PL are shown with blue, orange, and green lines, respectively. The error bar represents the standard error of the results of three equal-length AA MD blocks, each containing 1 µs.

**Figure 5.**
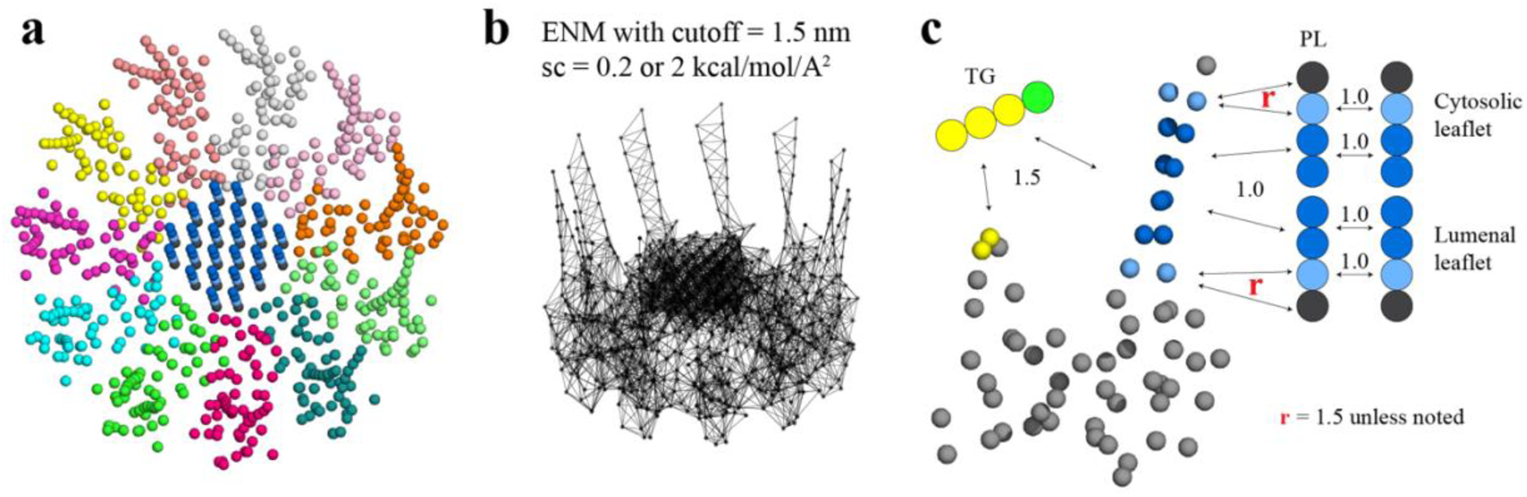
CG models of seipin and lipids. (a) CG model of human seipin oligomer. The CG atoms inside the hydrophobic helix ring represent PL atoms (proteinized PLs). (b) Elastic network model (ENM) with a spring constant of 0.2 or 2 kcal/mol/Å^2^. (c) Scaling factors of attraction parameters between seipin-PL and seipin-TG interactions. PL head, interfacial, and tail groups are shown with black, light blue, and dark blue, respectively. TG glycerol and tail groups are shown with green and yellow, respectively. Seipin atoms that attract PL tails are shown with dark blue, and those that attract PL interfacial atoms are shown with sky blue. Two seipin atoms in the HH, shown with yellow, and four central seipin atoms in each TM segment, shown with dark blue, attract TG atoms. Scaling factors (r) between seipin residues at the membrane interface and PL interfacial and head group are set to 1.5 unless noted.

LD biogenesis is a microscopic/mesoscopic process, having its time and length scales beyond those accessible by present day AA MD simulations. For instance, during our 3 µs-long AA MD simulation we observed recruitment of TG inside the seipin complex, but not TG nucleation. To access the relevant time and length scales, we instead developed CG lipid and seipin models (Fig. 5). Linear, four-site models were used for lipids (Grime and Madsen, 2019; Kim et al.). Every four protein residues were linearly mapped to one CG atom to match the resolution with CG lipids. We also placed 24 PL molecules inside the HH ring with the orientation consistent with other PL molecules in the lumenal leaflet because they were considered a part of a seipin oligomer. This is based on the AA MD simulation that demonstrated PLs inside the HH ring are trapped. We constructed an elastic network model (ENM) (Haliloglu et al., 1997) by connecting a pair of seipin CG atoms via an effective harmonic bond with a spring constant (sc) of 0.2 kcal/mol/Å^2^ if the distance is less than 1.5 nm (Fig. 5b). Such a choice was made to best reproduce the fluctuations from AA MD simulation data. To understand the impact of the stiffness of harmonic bonds, we also made a model with a spring constant of 2 kcal/mol/Å^2^. To achieve the known stability of seipin in a bilayer membrane, nonbonded protein-lipid interactions were based on the lipid-lipid interactions. Attraction scaling factors that change the force and potential depth linearly are shown in Fig. 5c (Kim et al.). Although it is not obvious how to quantitatively incorporate the AA MD simulation data into phenomenological CG models, higher scaling factors between TG-HH and TG-TM can be qualitatively justified by the high attractions of those pairs indicated in the analysis of the AA MD simulation (Fig. 2d).

Seipin is thought to catalyze TG nucleation, thereby decreasing the critical concentration (Chung et al., 2019). To test this hypothesis, we performed the CG simulations of spherical bilayers with a diameter of 40 nm. The curvature was comparable to the actual ER tubule (Georgiades et al., 2017). The initial structures had evenly distributed 2% TG molecules. Because the TG concentration is below the critical concentration for its phase transition (Hamilton and Small, 1981; Khandelia et al., 2010; Zoni et al., 2021a), TG nucleation did not occur in the lipid system (Fig. 6). In contrast, the system that includes the seipin complex showed a nucleated TG lens inside the complex due to the attractions between TG-TM and TG-HH (Fig. 6). As a control simulation, we included a single TM segment in the lipid system and carried out the CG MD simulation. TG nucleation did not happen in the system (Fig. 6), indicating that a single TM segment falls well short of TG nucleation. This result is consistent with the notion that not all TM proteins can facilitate TG nucleation despite that most hydrophobic residues in the PL tail region prefer interactions with TG than with PL, as shown in Fig. 2. Therefore, high protein density at the seipin site provides for collective and cooperative interactions with TG, catalyzing TG nucleation.

**Figure 6.**
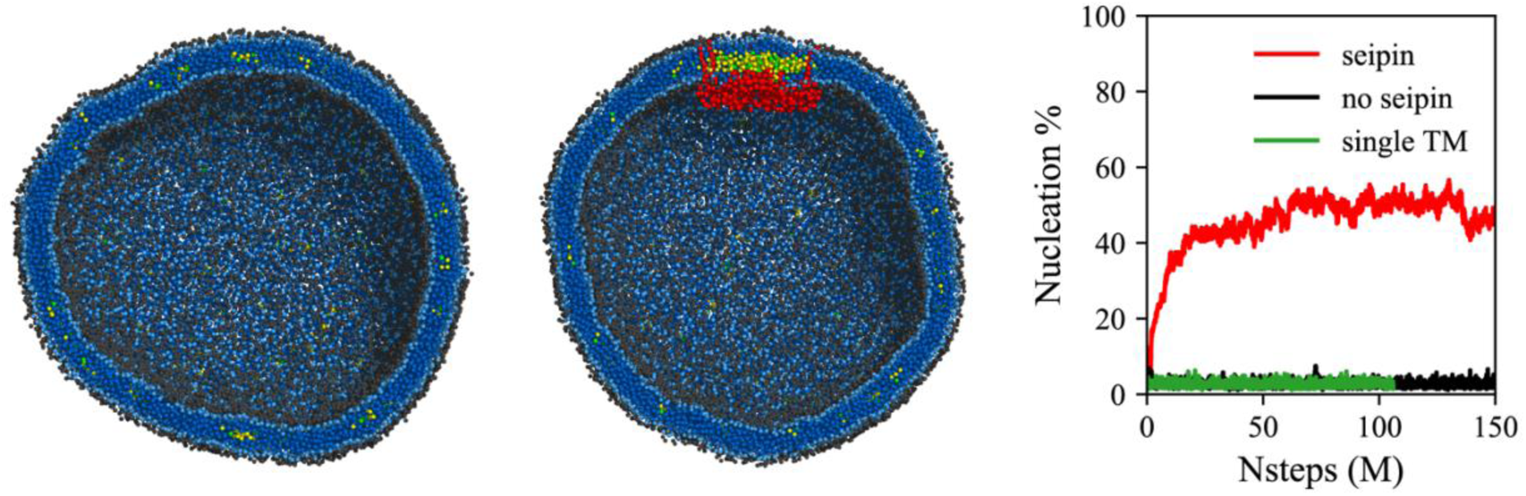
Seipin lowers the critical concentration of TG nucleation. CG MD simulations of bilayers containing 2% TG with a diameter of 40 nm were carried out. The clipped snapshots of the last frames of the pure lipid (left) and seipin-containing systems (center) are shown. The ENM model with a spring constant of 0.2 kcal/mol/Å^2^ was used. PL head, interfacial, and tail atoms are shown with black, light blue, and dark blue, respectively. TG glycerol and tail atoms are shown with green and yellow, respectively. Seipin oligomer is indicated with red. The nucleation percentages of three systems with simulation steps are shown in right.

To study the impact of the cage-like structure of the seipin oligomer and their TM segments in LD biogenesis, we simulated various geometries of seipin in the spherical bilayers containing 6% TG (Fig. 7). In the first model, we removed the TM segments, and the resulting model only contained the lumenal domain. In the second model, we removed 6 continuous subunits from the seipin oligomer, and the resulting model contained 5 subunits. As a reference, we also carried out simulations of the pure lipid system and the 11mer-containing system. The ENM with a spring constant of 0.2 kcal/mol/Å^2^ was used. Because the TG concentration was above the critical concentration, TG nucleation occurred in those systems even in the system without seipin. However, the resulting morphologies of oil lenses of those systems were different as shown in the final snapshots and characterized by anisotropy. First, in the lipid system, a nucleated TG lens was flat and had high anisotropy to minimize the membrane deformation penalty (Kim et al.). In contrast, in the 11-subunits model, a nucleated TG lens was located on top of the seipin lumenal domain, surrounded by seipin TM segments. This resulted in a significant change in the shape of the oil lens from high anisotropy in the lipid system, minimizing the membrane deformation penalty, to low anisotropy in the seipin-containing systems. Given the nucleation percentages were comparable in those simulations, a change in anisotropy can be attributed to the seipin TM, not to the amount of nucleated TG molecules. Importantly, the seipin TM segments constrained the XY area where TG can be in the bilayer, pushing excessive TG molecules to the budding LD. This results in the formation of the ER-LD neck structure, consistent with an experimental tomogram (Salo et al., 2019). An equilibrated diameter of the ER-LD structure (Fig. S6), approximated by a diameter of the circle formed by the end residues of N- and C-terminal TM segments, also agreed well with the experimentally measured data, which is in the range of 13 to 17 nm (Salo et al., 2019). We note the TM segments tilted away from the oligomeric center during LD growth; therefore, the diameter of the seipin ring increased with simulation time. We also simulated the 11-subunit model with a spring constant of 2.0 kcal/mol/Å^2^ (Fig. S7). The higher spring constant reduced the diameter of the ER-LD neck structure. However, the nucleation percentage and morphology of the formed oil lens had little difference with the previous system that contained the ENM with a spring constant of 0.2 kcal/mol/Å^2^.

**Figure 7.**
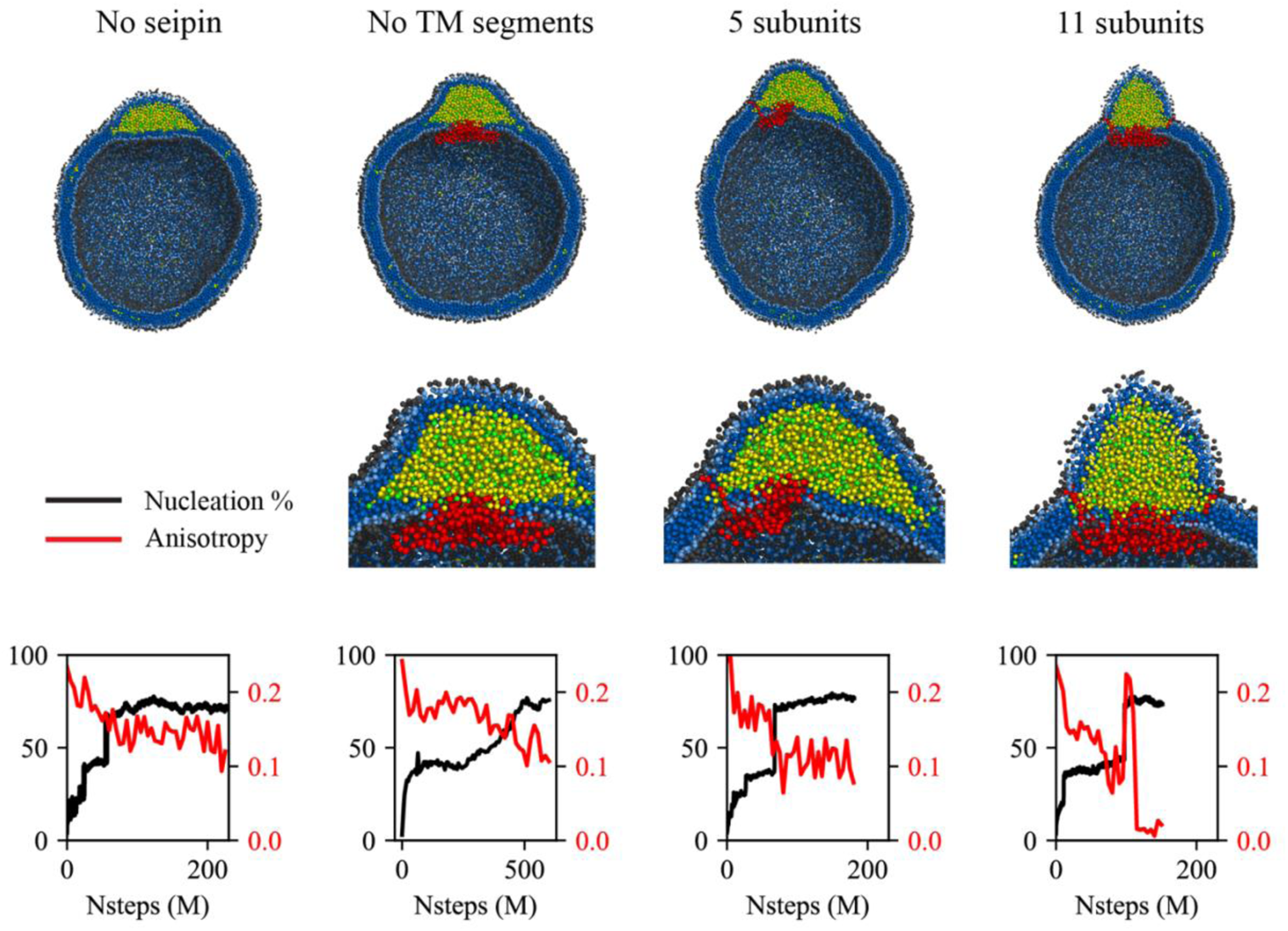
Cage-like geometry and neck formed by seipin TM segments are key to modulating the morphology of a forming oil lens. The first row shows the clipped snapshots of the equilibrated frames, and the second row shows the close-up view of the seipin. The color code is as in Fig. 6. The third row shows the nucleation percentage (black) and anisotropy (red). CG simulations of spherical bilayers containing 6% TG with a diameter of 40 nm were carried out. The ENM model for seipin with a spring constant of 0.2 kcal/mol/Å^2^ was used.

The 5-subunit model can be considered a mixture of the lipid system and 11-subunit model because one end is occupied with seipin subunits while the other end is exposed to lipids. The resulting morphology of an oil lens from the CG MD simulations was also between those results. The TG oil lens was elongated to the region where there was no seipin subunit; however, it was constrained in the region of seipin subunits, especially by their TM segments. The equilibrated anisotropy was also between the lipid and 11-subunits systems. Finally, we simulated the seipin model that only contained the lumenal domain. Such a complex does not form a mobile focus in cells likely because it fails to form an oligomer, it is not stable in bilayers, or it is degraded (Chung et al., 2019). However, simulating this system can provide further valuable insight on the roles of the TM segments. The resulting oil lens showed little difference from the lipid system without seipin. The anisotropy was high, and the formation of the ER-LD neck structure was abolished.

It is worth noting that the mechanisms of oil growth can be predicted from analysis of the nucleation percentage and anisotropy. A sharp increase in the nucleation percentage indicates oil coalescence, as shown in ~70 M MD time steps in the 5 subunit system and ~100 M MD time steps in the 11 subunit system. In contrast, the nucleation percentage grew slowly in the seipin model without the TM segments from ~200 M MD time steps, indicating Ostwald ripening. Finally, a sudden increase followed by a sharp decrease in anisotropy in the 11 subunits system implies a slow coalescence. When any two TG molecules in distinct oil lenses are within 2 nm, those oil lenses are considered one oil lens, as shown by a step increase in the nucleation percentage at 100 M MD time steps. Such a snapshot resembles two humps, increasing anisotropy. After ~110 M MD time steps, the oil lenses are fully merged into one spherical lens, reducing anisotropy to zero. Controlled coalescence at the seipin site will be discussed later.

To investigate the more advanced biogenesis steps, we simulated a larger spherical bilayer with a diameter of 60 nm and 6% TG. We also constructed a hENM model (Lyman et al., 2008) using the fluctuations obtained from the AA simulation of seipin in the bilayer membrane (Fig. 8a). The calculated RMSF from the AA and CG simulations agreed well in bilayers (Fig. 8b). Consistent with the previous results, the seipin TM segments defined the oil boundary, facilitating the transport of TG into the LD (Fig. 8c). The equilibrated anisotropy was close to zero, indicating a spherical shape of the forming oil lens (Fig. 8d). Collectively, our tests demonstrate that the ring geometry of seipin TM segments is key to defining the boundary of the forming oil lens and creating the unique ER-LD structure.

**Figure 8.**
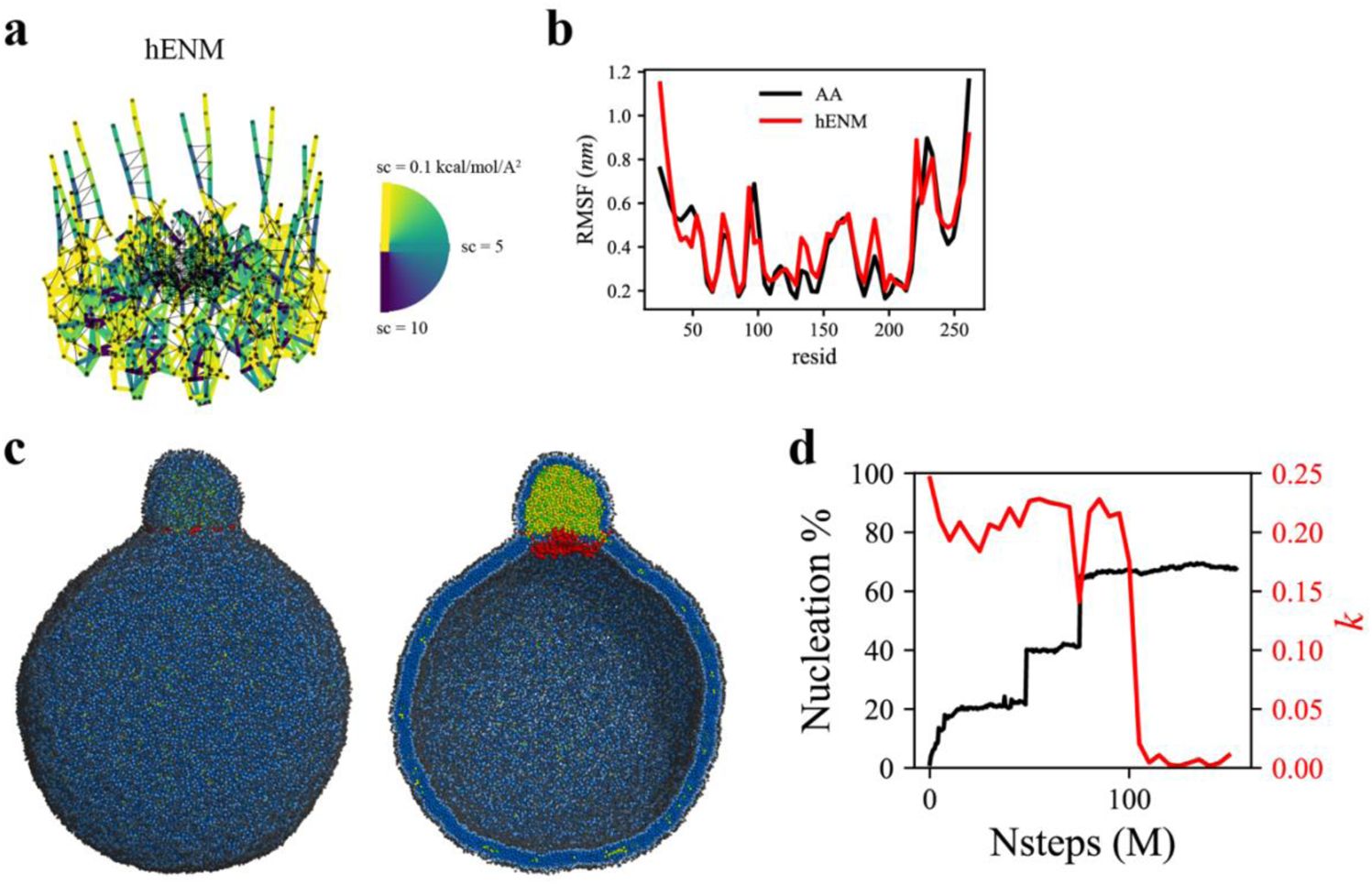
CG-MD shows LD growth in a large bilayer. CG MD simulations of spherical bilayers containing 6% TG with a diameter of 60 nm were carried out. (a) Heterogeneous ENM (hENM) model of human seipin was constructed. Pairs of atoms in a subunit were connected with harmonic springs with their spring constants (sc) represented by their color. Additional harmonic springs (black lines) with a spring constant of 0.1 kcal/mol/Å^2^ were added between pairs of atoms that were not included in the hENM with a distance cutoff of 11 Å to ensure connections between subunits. (b) RMSF of a seipin subunit was compared between the AA trajectory and hENM in a bilayer. (c) Exterior and interior view of the last frame of the CG simulation. The color code is as in Fig. 6. (d) The nucleation percentage (black line) and anisotropy (red line) are shown.

The coevolutionary sequence analysis (Hopf et al., 2019) of human seipin indicated that seipin’s lumenal domain and two TM segments are evolutionarily conserved (Cartwright and Goodman, 2012) (Fig. S8). In contrast, the N- and C-terminal tail regions, exposed to the cytosol, are not conserved (Fig. S8). To test whether the non-conserved, cytosolic tails are important for LD formation, we experimentally constructed seipin-ΔTERM, which lacks the N-terminal region (1-22 amino acids) and C-terminal region (268-398 amino acids). First, we confirmed that seipin-ΔTERM assembles discrete seipin foci in the ER, the same as seipin full-length (FL) protein. We next tested whether seipin-ΔTERM is functional by examining its ability to rescue the LD phenotypes in seipin knockout (KO) SUM159 cell line we previously reported (Wang et al., 2016). Consistent with the previous experiments (Chung et al., 2019; Fei et al., 2008; Salo et al., 2016; Szymanski et al., 2007; Wang et al., 2016), seipin KO cells had massive accumulation of small nascent LDs at the early time (~1 hours) of LD formation after oleate treatment (Fig. 9a). Seipin-ΔTERM rescued the defective LD phenotype of seipin knockout cells (Fig. 9a), indicating that the cytosolic non-conserved N-/C-terminal regions of human seipin are dispensable for the seipin function. This is comparable with previous experiments that demonstrated the removal of N- or C-terminal region of fly seipin in *Drosophila* S2 cells did not change the rescue efficiency of the seipin deletion phenotype (Wang et al., 2016).

**Figure 9.**
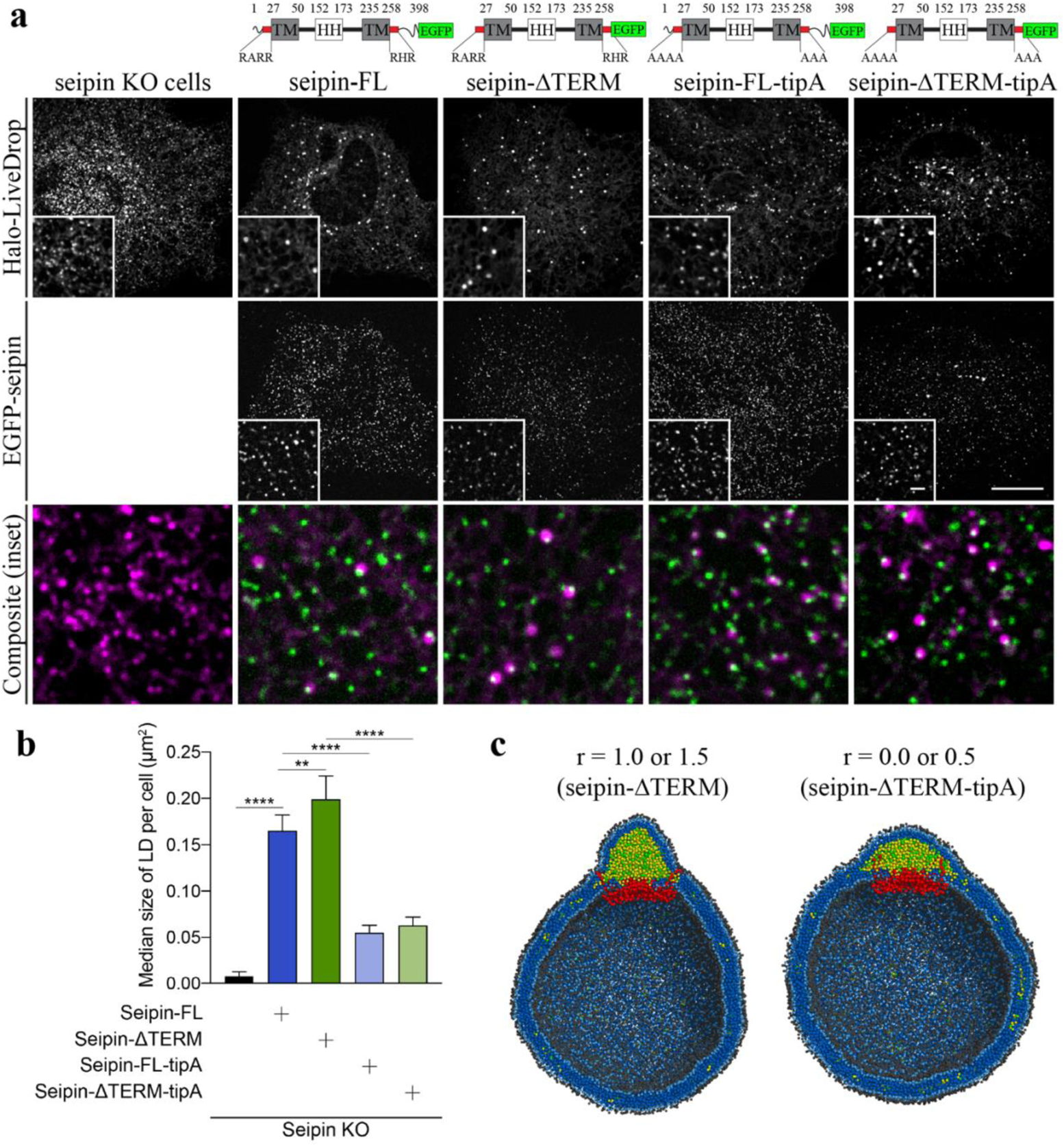
Non-conserved, cytosolic tails of human seipin are dispensable for the function, while the conserved, positively charged residues at the ends of seipin TM segments are critical for LD maturation. (a) Confocal imaging of live seipin KO SUM159 cells transiently transfected with various seipin constructs fused with EGFP and Halo-LiveDrop (stained with JF549). The cells were pre-incubated with 0.5 mM oleic acid for 1 h prior to image acquisition. Scale bars, full-size, 15 μm; inserts, 2 μm. (b) Quantification of size of LDs per cell shown in (a). n = 4 cells. More than 300 LDs were analyzed in each sample. Median with interquartile range. \*\*\*\**p* < 0.0001, \*\**p* < 0.01 were calculated by unpaired t test. (c) CG MD simulations with various attraction scaling factors (*r*). The spherical bilayer has a diameter of 40 nm and contains 6% mol TG. The ENM model of human seipin with a spring constant of 0.2 kcal/mol/Å^2^ was used. The color code is as in Fig. 6.

We also investigated if the positively charged residues located at the beginning of the N-terminal TM segment (R23, R24, R26) and the end of the C-terminal TM segment (R265, H266, R267) are essential for the seipin function. Those charged residues located at the borders of the TM segments are conserved (Fig. S8), and in our CG trajectories, they were exposed to the outer leaflet of the membrane (Figs. 7 and 8c). To test the importance of those residues, we experimentally made the two mutant seipin constructs, seipin-FL-tipA and seipin-ΔTERM-tipA, in which their arginine and histidine residues at the ends of seipin’s TM segments were mutated to alanine. Despite the absence of charged residues, those seipin constructs stably formed foci at the ER bilayer and resulted in an intermediate LD phenotype with more and irregular shaped LDs than are formed in wild type cells (Figs. 9a and 9b).

We further hypothesized that the reduced attraction between the borders of seipin’s TM segments and the membrane interface caused defective LD maturation in the cells that contained the mutant constructs. To investigate this possibility, we carried out CG simulations with variable attraction scaling factors (*r*) between the residues at the membrane interface and PL interfacial and head groups (Figs. 5c and 9c). When *r* is small, it could be thought of as the mutant construct that does not have the charged residues at the ends of TM segments (seipin-ΔTERM-tipA). The resulting CG simulations demonstrated that those residues no longer interacted with the interface of the cytosolic leaflet (Fig. 9c). Instead, the whole TM segments were immersed in the forming oil lens, leading to the destruction of the ER-LD neck structure. In particular, the residues at the end of TM segments lost contact with the outer membrane during oil coalescence when a seipin-free oil lens approached the seipin ring and merged with the oil lens contained within it. The loss of the ER-LD neck structure resulted in a flat oil lens, as shown in our CG system that did not include seipin or that had seipin without TM segments (Fig. 7). In contrast, the ER-LD neck structure was maintained during oil coalescence when *r* was large, as in the previous CG simulations. TG molecules in a seipin-free oil lens were transferred to the seipin in a controlled manner without disrupting the ER-LD neck structure. Consistent with our previous CG results (Figs. 7 and 8), the seipin TM segments served as a boundary between the ER bilayer and the resulting oil lens, creating the ER-LD neck structure. Collectively, the cell experiments and CG simulations suggest that the conserved, charged residues located at the borders of seipin TM segments maintain the ER-LD neck structure during oil coalescence and promote LD growth.

## DISCUSSION AND CONCLUSIONS

Seipin is a critical protein that orchestrates LD formation (Arlt et al., 2021; Chung et al., 2019; Prasanna et al., 2021; Salo et al., 2020; Salo et al., 2019; Wang et al., 2016; Zoni et al., 2021b). Recent joint computational and experimental studies reported that the HH of seipin facilitates TG nucleation (Prasanna et al., 2021; Zoni et al., 2021b). However, little is known about the roles of seipin TM segments despite their experimentally confirmed importance for seipin’s function (Arlt et al., 2021; Chung et al., 2019). Capitalizing on highly CG models of lipids and seipin, we show that a cage-like geometry of seipin TM segments forms a constricted neck, converting a planar oil lens into a unique ER-LD, facilitating LD growth. In contrast, the system that lacked seipin or contained seipin without TM segments resulted in a flat oil lens with high anisotropy.

Using cell-based experiments and CG simulations, we further identified certain essential and dispensable parts of human seipin. We show that the non-conserved N- and C-terminal cytosolic regions of human seipin are not required for LD formation. Therefore, truncated seipin models used in the current and previous computational studies are reasonable due to the dispensability of the cytosolic tails (Prasanna et al., 2021; Zoni et al., 2021b). We also provide evidence that the conserved, positively charged residues located at the borders of the seipin TM segments are crucial for LD maturation. Mutating those residues to alanine resulted in an intermediate LD phenotype with more and nonuniform shaped LDs than those formed in wild type cells. In the CG simulations, seipin TM segments were immersed in an oil lens when interactions between those residues and PL interfacial and head groups became reduced.

Based on our data, we propose a model in which the positively charged residues, located at the borders of seipin TM segments, anchor the TM segments at the cytosolic side of the membrane (Fig. 10). This was particularly important in maintaining the ER-LD neck structure during oil coalescence. When a seipin-free oil lens approaches a seipin-positioned oil lens, the TM segments should keep their positions in the bilayer to maintain the ER-LD neck structure. Strong electrostatic interactions between the seipin residues at the borders of seipin TM segments and the membrane interface of the cytosolic leaflet inhibited rapid coalescence. Instead, TG in the seipin-free lens was transported to the seipin-positioned oil lens in a controlled manner. A slow coalescence at the seipin site is implicated in anisotropy analysis in Fig. 7. If the interactions between the seipin TM tip residues and the cytosolic leaflet are small, the seipin TM segments become immersed in the lens during swift coalescence, affecting LD maturation.

**Figure 10.**
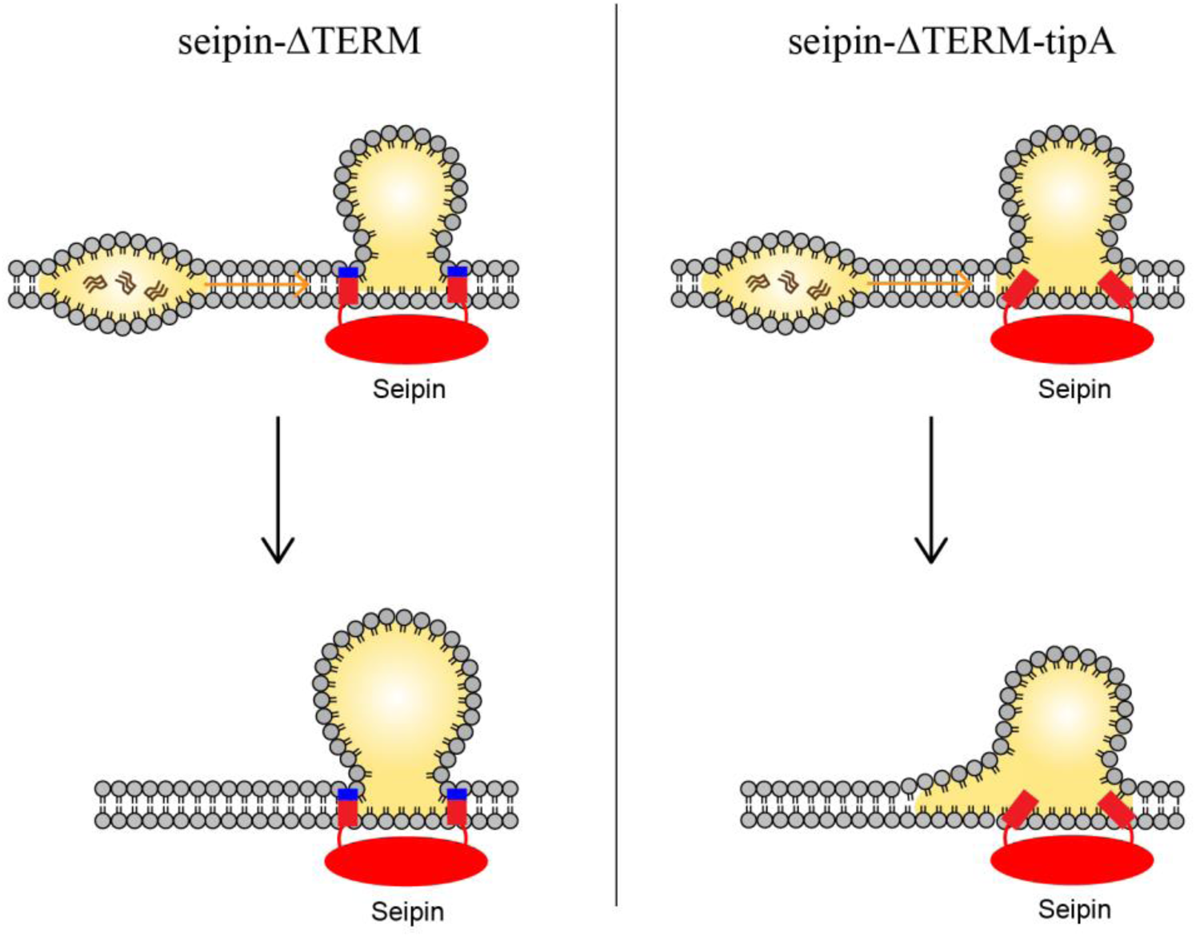
Model for the role of the charged residues at the end of seipin TM segments during oil coalescence. The left side shows the seipin TM segments maintain the bilayer thickness around the ER-LD neck structure. The right side shows the TM segments are immersed in the oil lens during oil coalescence because positively charged residues at the end of seipin TM segments (blue) are mutated to alanine.

We investigated protein-lipid interactions and lipid dynamics using the AA simulation. Hydrophobic interactions between seipin TM segments and HH with TG are the main driving force of TG nucleation, in conjunction with hydrophilic interactions between the TG glycerol moiety and protein residues. It is worth noting that different force fields consistently report a strong attraction between S166 and TG glycerol moieties (Prasanna et al., 2021; Zoni et al., 2021b). During LD growth, lipids or proteins such as *LiveDrop* migrate from the ER onto LD through the populated TM region (Olarte et al., 2020; Wang et al., 2016). We speculate that the collective motions of TM segments will increase the permeation of proteins and lipids by opening the space between neighboring monomers. Also, as demonstrated in the CG simulations, the diameter of the ER-LD neck structure during the LD growth phase is larger than during the initial nucleation stage. The widely spread TM segments will promote lipid and protein influx to LDs.

Finally, we discuss the limitations of our study. First, LDAF1 is not included in our simulations as its structure and number in the seipin oligomer are not identified yet. LDAF1 is predicted to have a double hairpin structure, which increases the density of TM segments inside the seipin oligomer. Therefore, including LDAF1 in simulations will likely change protein-lipid interactions in the system and lipid diffusion. Second, the CG simulations benefited from highly CG models to simulate large systems and study advanced LD biogenesis steps. However, seipin sequence-specific features are missing in the seipin model. Also, since this is a phenomenological model, the quantities calculated are not directly related to the underlying AA systems.

Taken together, we suggest a model in which seipin TM segments are key for LD biogenesis by nucleating TG, controlling lipid diffusion, defining the boundary of the forming oil lens, maintaining the ER-LD neck structure, and controlling oil coalescence. Our study thus provides a broader and deeper understanding of the roles of the TM segments, critical for seipin function.

## MATERIALS AND METHODS

### Seipin structure

We modeled the seipin structure (Arg23-Arg265) based on our cryoelectron microscopy data (Fig. S1). Our structure contained the lumenal domain (Val60-His219), which was previously resolved (Chung et al., 2019; Yan et al., 2018), and the partially resolved TM segments. However, due to the low resolution of the TM region, we were not able to identify residues in the TM segments. Instead, we used the orientation of the TM helices in our modeling. We assumed the residues that corresponded to the N- and C-terminal TM helical structures were Leu29-Gly50 and Ala235-Val258, respectively. The missing residues from Ser51 to Val60 were determined with Modeller (Fiser et al., 2000). The structure from Phe220 to Phe230, referred to as a switch region, was homology modeled using a yeast seipin structure as a reference because this region is highly conserved and predicted to be folded similarly (Arlt et al., 2021). The structure of the switch region was helical, and its helicity was further supported by the PSIPRED (McGuffin et al., 2000), TMHMM (Krogh et al., 2001), TMpred, and Phyre2 (Kelley et al., 2015) servers. The missing residues, Pro231-Cys234, were modeled with Modeller (Fiser et al., 2000).

### All-atom (AA) MD simulation

The seipin simulation in a POPC bilayer including 6% TG was carried out for 3 µs. The equilibrated bilayer structure was taken from the previous work (Kim and Voth, 2021). Seipin has a HH ring at the center with a radius of ~2 nm in the lumenal leaflet. We first placed 20 POPC molecules inside the seipin HH ring with their orientations consistent with other lumenal POPC molecules using PACKMOL (Martínez et al., 2009). Those PLs remained trapped inside the ring during the simulation (proteinized PLs). We put the seipin complex at the membrane center and removed any lipid molecules within 0.9 Å of seipin atoms. The equilibrium protocol provided by the CHARMM-GUI interface was used (Lee et al., 2016). Additionally, 100 ns of equilibration was carried out, restraining the positions of the backbone atoms of the lumenal domain (Val60-His219) and the Z positions of phosphorus atoms with a spring constant of 20 kJ/mol/nm^2^. The total numbers of POPC and TG molecules were 797 and 48, respectively.

The simulation was run by GROMACS 2020 (Van Der Spoel et al., 2005) with a Lennard-Jones (LJ) cutoff-free version of C36 (Yu et al., 2021a; Yu et al., 2021b). The modified TG parameters that reproduced the interfacial tension against water were used (Kim and Voth, 2021). Simulations were evolved with a 2-fs timestep. The long-range electrostatic and LJ interactions were evaluated with the Particle Mesh Ewald algorithm, with the real-space cutoff distance of 1.0 nm (Essmann et al., 1995). Bond involving a hydrogen atom was constrained using the LINCS algorithm (Hess, 2008). A temperature of 310 K and a pressure of 1 bar were maintained with the Nose-Hoover thermostat and the Parrinello-Rahman barostat, respectively (Hoover, 1985; Nosé, 1984; Parrinello and Rahman, 1981). The coupling time constants of 1 ps and 5 ps were used, respectively. A compressibility of 4.5 × 10^−5^ bar^−1^ was used for semi-isotropic pressure coupling.

### Coordination number analysis

To study protein-lipid interactions, we first reduced the resolution of the AA simulation by mapping each POPC molecule into 11 molecular groups and each TG molecule into 13 groups. In this mapping scheme, each POPC molecule had the choline head group, phosphate group, glycerol moiety and four tail groups for each acyl chain. Similarly, each TG molecule had the glycerol moiety and four tail groups for each acyl chain. Each protein residue was mapped into one backbone and one side chain group. For each amino acid, the coordination number between the side chain group and membrane groups of PL or TG was calculated by *S*_*M*_ = ∑_*M*_ ∑_*a*_[1 − (*r*_*a*_/*r*_0_)^6^] / [1 − (*r*_*a*_/*r*_0_)^12^], where *M* represents PL or TG and *a* represents a CG atom belonged to *M*. The parameter *r*_0_ was set to 0.4 nm and *r*_*a*_ is the distance between the side chain group and membrane group (atom *a*). The normalized coordination number by molecule or the coordination number per molecule (||*S*||) was computed by diving the coordination number by the number of molecules of PL or TG. The normalized coordination number by CG atom or the coordination number per CG atom (||*S*_*A*_||) was calculated by dividing the coordination number by the number of atoms of atom *A*.

### Principal component analysis (PCA) and diffusion coefficient

Using the AA trajectory, we carried out the PCA and diffusion coefficient calculation with the MDAnalysis library (de Buyl, 2018; Michaud-Agrawal et al., 2011). For the PCA, each monomer trajectory was globally aligned with the initial monomer structure. The coordinates of the α-carbon atoms were extracted from the aligned trajectory and used to compute the covariance matrix. The two primary collective motions with the highest explained variances were calculated. When calculating diffusion coefficients, we translated the system such that the center of the mass of the lumenal domain of seipin was at the origin in each frame. Therefore, the diffusion coefficient of PL or TG reported here represents the relative diffusion coefficient with respect to the center of mass of the protein. The trajectory was divided into three trajectories, each of which was 1 µs-long. In each trajectory, PL or TG molecules were categorized into three classes, based on the average XY-distance from the origin. The first class of lipids were located within 3.5 nm from the origin, slightly greater than the radius of the HH ring. The second class of lipids were located between 3.5 nm and 7.0 nm from the origin, where they mainly interacted with the TM segments. Finally, the lipids that were further than 7.0 nm were considered lipids in the protein-free zone as they did not interact with the protein. The position-dependent diffusion coefficients were reported by calculating diffusion coefficients in each class. Three equal-length blocks were used to report the average and standard errors.

### Coarse-grained (CG) simulation

We used a previously developed CG model for PL and TG with each molecule consisting of 4 CG beads (Grime and Madsen, 2019; Kim et al.). An angle parameter of 0.5 *k*_B_*T* for PL was used in this study. A CG model for seipin was constructed by linearly mapping four amino acids into each CG bead. Such a resolution was chosen to match the resolution of the CG lipids, preventing hydrophobic mismatch. We also placed 24 PL molecules inside the HH ring with their orientations consistent with the other PL molecules in the lumenal leaflet. Three models were constructed with different elastic networks with bond potentials *k*(*r* − *r*_0_)^2^ where *k* is the spring constant and *r*_0_ is the equilibrium bond length. The first two models utilized a standard ENM with a spring constant of 0.2 kcal/mol Å^2^ or 2 kcal/mol Å^2^ and with a distance cutoff of 15 Å. The third model utilized the hENM that more correctly represented the fluctuations of seipin in the underlying AA simulation (Lyman et al., 2008). From the AA MD simulation, we first made a concatenated, aligned seipin monomer trajectory and obtained the hENM with a cutoff distance of 12 Å. The hENM was applied to each monomer. A spring constant of 0.1 kcal/mol Å^2^ was applied to the CG pairs that did not have hENM if the distance is less than 11 Å. To achieve the known stability of seipin in a bilayer membrane, nonbonded protein-lipid interactions were based on the lipid-lipid interactions (Grime and Madsen, 2019; Kim et al.). Protein atoms located in the hydrophobic phase interacted with the PL tail atoms with the equal attraction strength of the pair between PL tail and PL tail atoms or PL interfacial and PL interfacial atoms (scaling factor = 1). Protein atoms located at the membrane interface attracted PL interfacial atoms with a scaling factor of 1.5 unless otherwise noted. The central region of TM segments and two HH beads attracted TG atoms with a scaling factor of 1.5. Every CG bead carried a mass of 200 g/mol and no charge. Spherical bilayers with a diameter of 40 nm or 60 nm were constructed, containing 2% TG or 6% TG, followed by the placement of seipin and removal of lipids that had a close contact with seipin.

The CG simulations were with by LAMMPS MD software with tabulated CG potentials (Plimpton, 1995). Simulations were evolved with a 50-fs timestep. Temperature was maintained at 310 K by the Langevin thermostat with a coupling constant of 100 ps (Schneider and Stoll, 1978). The cutoff distance of nonbonded interaction was 1.5 nm. The initial structures of CG simulations were prepared with the MDAnalysis library (Michaud-Agrawal et al., 2011).

### Nucleation percentage and anisotropy

We calculated the nucleation percentage and anisotropy as explained in (Kim et al.). In short, the nucleation percentage was defined as the ratio of the number of TG molecules in the largest cluster to the number of TG molecules in the system. The distance cutoff of 2 nm was used for clustering TG molecules. The anisotropy was calculated by diagonalizing the moment of inertia tensor of the largest TG cluster. The anisotropy of 0 represents a spherical shape, and the anisotropy of 0.25 represents a planar shape.

### Plasmid Construction

For plasmid construction, all PCRs were performed using PfuUltra II Fusion HotStart DNA polymerase (#600672, Agilent Technologies) and restriction enzymes were from New England Biolabs. The fragment DNA of tipA mutant construct was synthesized (gBlock, Integrated DNA Technologies).

### Fluorescence Microscopy

Cells were plated on 35 mm glass-bottom dishes (MatTek Corp). Imaging was carried out at 37°C approximately 24 h after transection. Before imaging, cells were transferred to pre-warmed FluoroBrite DMEM supplemented with 2 mM GlutaMAX (#35050061, Thermo Fisher Scientific), 5 mg/mL insulin (Cell Applications), 1 mg/mL hydrocortisone (Sigma Aldrich), 5% FBS (Life Technologies 10082147; Thermo Fisher), 50 mg/mL streptomycin, and 50 U/mL penicillin.

Spinning-disk confocal microscopy was performed using a Nikon Eclipse Ti inverted microscope equipped with Perfect Focus, a CSU-X1 spinning disk confocal head (Yokogawa), Zyla 4.2 Plus scientific complementary metal-oxide semiconductor (sCMOS) cameras (Andor, Belfast, UK), and controlled by NIS-Elements software (Nikon). To maintain 85% humidity, 37°C and 5% CO_2_ levels, a stage top chamber was used (Okolab). Images were acquired through a 603 Plan Apo 1.40 NA objective or 1003 Plan Apo 1.40 NA objective (Nikon). Image pixel size was 0.065 mm. Green or red fluorescence were excited by 488 or 560 nm (solid state; Andor, Andor, Cobolt, Coherent, respectively) lasers. All laser lines shared a quad-pass dichroic beamsplitter (Di01-T405/488/568/647, Semrock). Green and red emission was selected with FF03-525/ 50 or FF01-607/36 filters (Semrock) respectively, mounted in an external filter wheel. Multicolor images were acquired sequentially.

### Imaging Quantification

LD size quantification was done with the fiji software. Individual images were first converted to a binary mask using “Threshold” plugin in fiji with Otsu method. Then, LD sizes were measured by the fiji plugin “Particle Analysis”.

## ACKNOWLEDGMENTS

This research was supported by National Institutes of Health (NIH) grants R01-GM063796 (to S.K. and G.A.V.), NIH R01-GM124348 (to R.V.F.), and NIH R01-GM097194 (to T.C.W.). The computer simulations were performed on the Stampede2 supercomputer at the Texas Advanced Computing Center and the Bridges2 supercomputer at the Pittsburgh Supercomputing Center (PSC) through allocation MCA94P017 with resources provided by the Extreme Science and Engineering Discovery Environment (XSEDE) supported by NSF grant ACI-1548562. We also utilized computational resources on the Midway3 supercomputer at the University of Chicago. J.C. is a fellow of the Damon Runyon Cancer Research Foundation. T.C.W. is a Howard Hughes Medical Institute Investigator. We thank Xudong Wu for performing cryo-electron microscopy imaging and critical discussion. S.K. acknowledges Chenghan Li and Sriramvignesh Mani for critical discussion.

## Supporting Information

**Figure S1.**
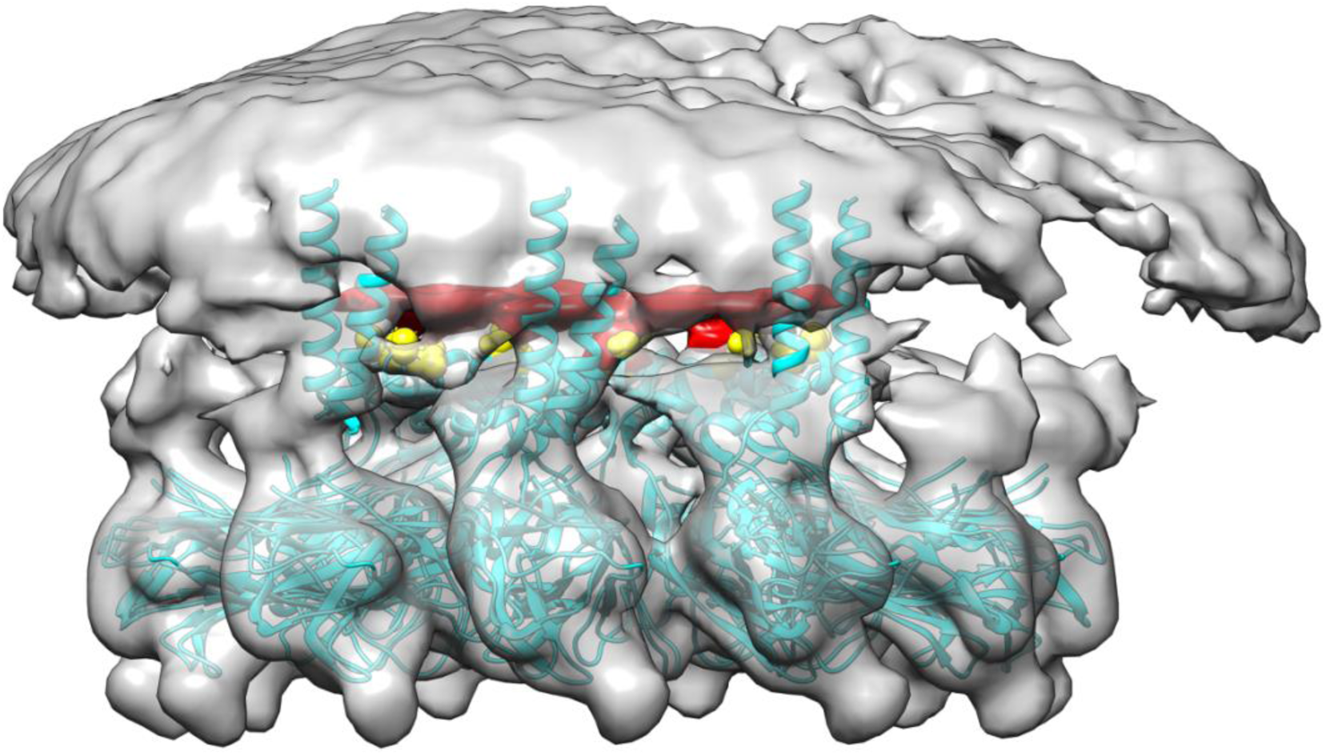
Cryoelectron microscopy of human seipin. The cyan model includes the seipin luminal domain and partially resolved transmembrane segments. The electron density of Ser165 and Ser166 is shown with yellow. The unidentified density that interacts with Ser165 and Ser166 is shown with red.

**Figure S2.**
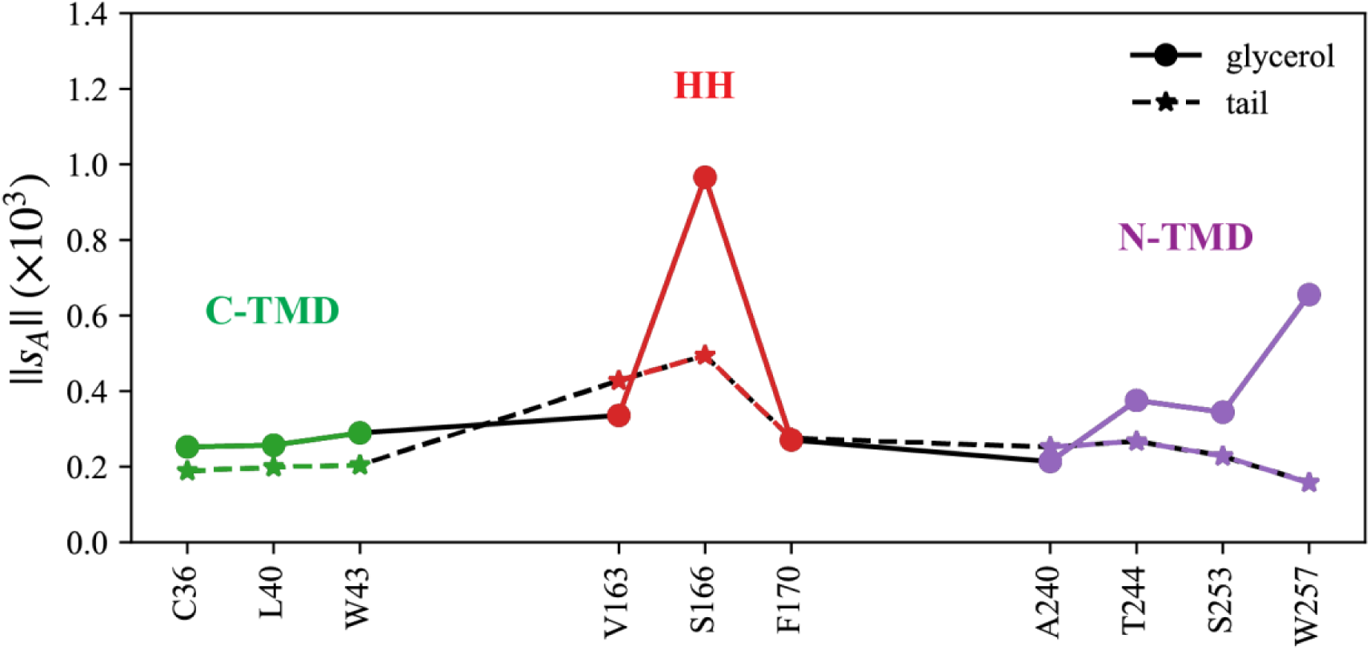
Normalized coordination number by atom. The interactions with TG glycerol moiety are shown with a continuous line and circle markers and those with TG tail atoms are shown with a dashed line and star markers. Related to Fig. 2 of the main text.

**Figure S3.**
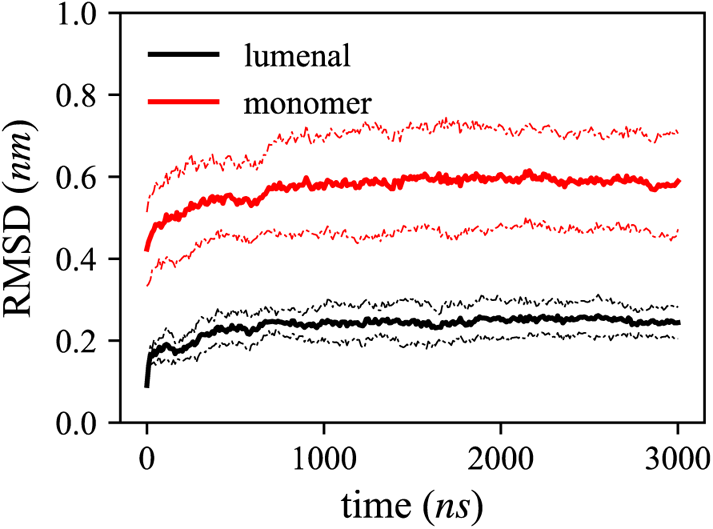
Root-mean-square distance (RMSD) of the α-carbon atoms of the lumenal domain (black) or the whole subunit (red). The error bars represent the standard deviation of the RMSDs of 11 subunits.

**Figure S4.**
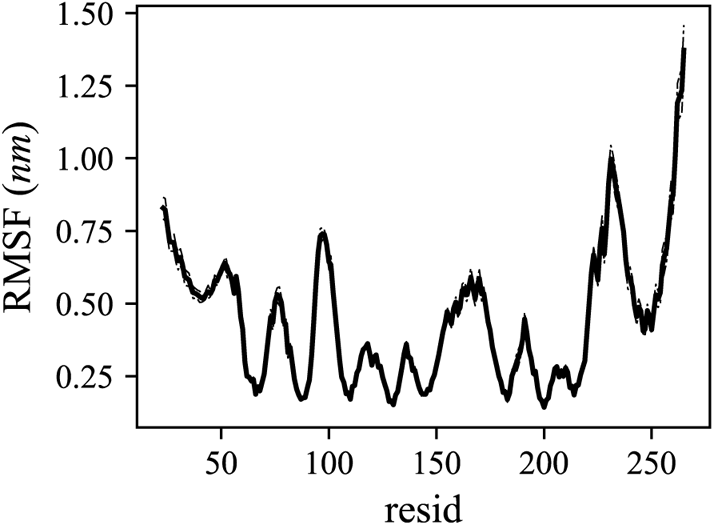
Root-mean-square fluctuation (RMSF) of the α-carbon atoms. The error bars represent the standard error of the RMSFs of three blocks.

**Figure S5.**
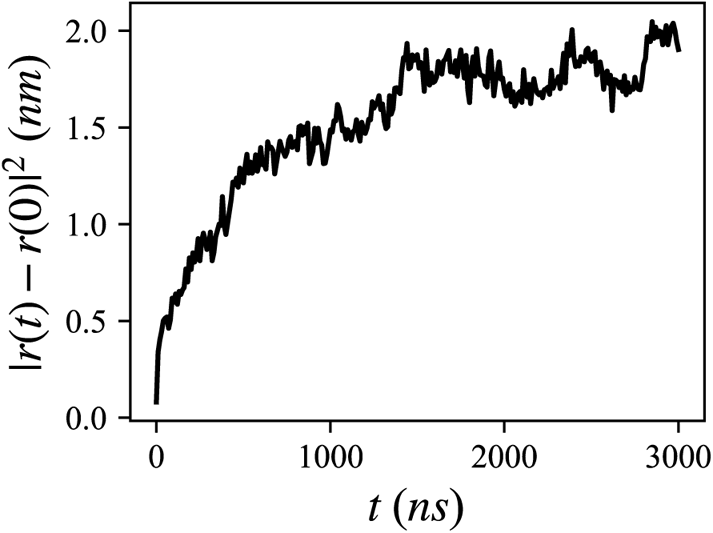
Mean squared distance of the 20 lumenal PLs, trapped inside the hydrophobic helix.

**Figure S6.**
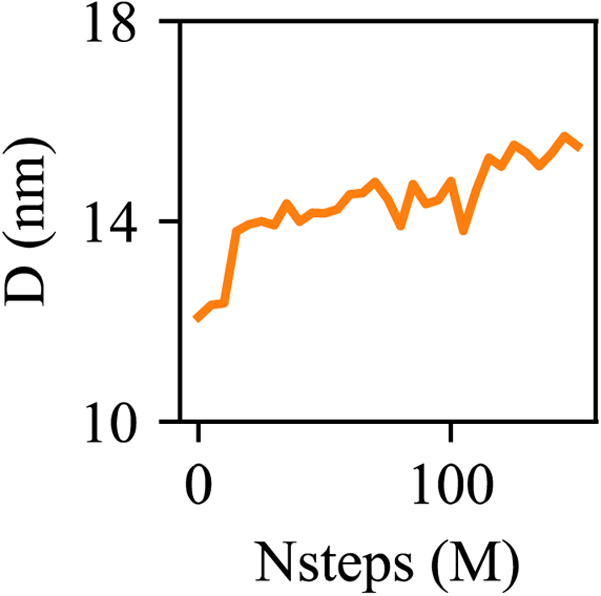
The diameter of an oil lens is shown with simulation steps.

**Figure S7.**
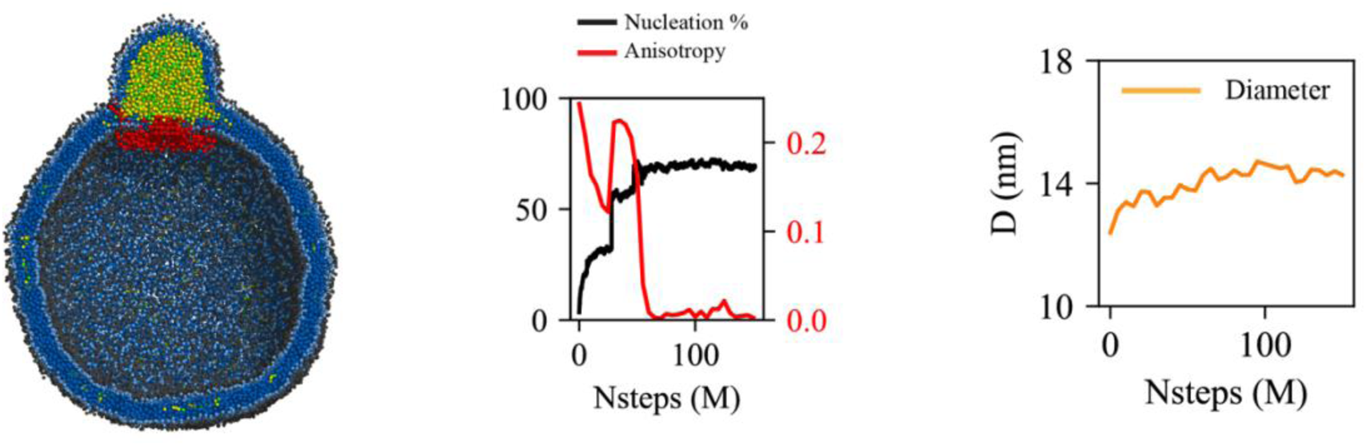
CG simulation of the bilayer containing 6% TG with a diameter of 40 nm. The ENM of seipin used a spring constant of 2 kcal/mol/Å^2^. The clipped snapshot is shown in left, the nucleation percentage (black) and anisotropy (red) are shown in middle, and the diameter of an oil lens (orange) is shown in right.

**Figure S8.**
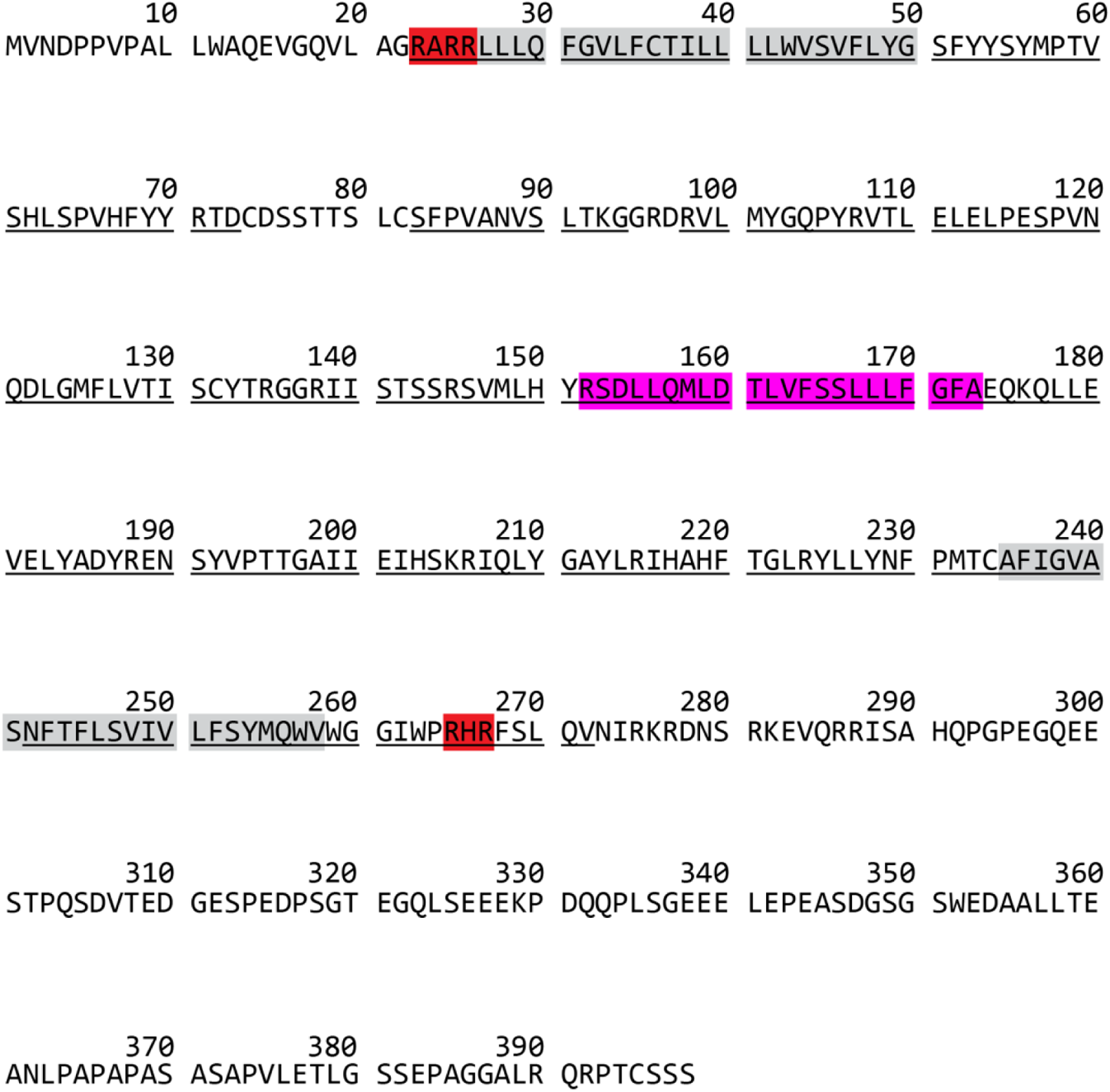
Sequence of human seipin. The conserved residues were underlined. Residues highlighted with red are positively charged residues, located at the seipin TM tips. Residues highlighted with gray or pink represent the TM segments or hydrophobic helix, respectively.

